# Human Anterior Insular Cortex Encodes Multiple Electrophysiological Representations of Anxiety-Related Behaviors

**DOI:** 10.1101/2024.03.05.583610

**Authors:** A Moses Lee, Virginia E Sturm, Heather Dawes, Andrew D Krystal, Edward F Chang

**Author notes:** Corresponding Author: A Moses Lee, MD PhD University of California, San Francisco Department of Psychiatry and Behavioral Sciences Weill Institute for Neurosciences 401 Parnassus Ave San Francisco, CA 94158.

## Abstract

Anxiety is a common symptom across psychiatric disorders, but the neurophysiological underpinnings of these symptoms remain unclear. This knowledge gap has prevented the development of circuit-based treatments that can target the neural substrates underlying anxiety. Here, we conducted an electrophysiological mapping study to identify neurophysiological activity associated with self-reported state anxiety in 17 subjects implanted with intracranial electrodes for seizure localization. Participants had baseline anxiety traits ranging from minimal to severe. Subjects volunteered to participate in an anxiety induction task in which they were temporarily exposed to the threat of unpredictable shock during intracranial recordings. We found that anterior insular beta oscillatory activity was selectively elevated during epochs when unpredictable aversive stimuli were being delivered, and this enhancement in insular beta was correlated with increases in self-reported anxiety. Beta oscillatory activity within the frontoinsular region was also evoked selectively by cues-predictive of threat, but not safety cues. Anterior insular gamma responses were less selective than gamma, strongly evoked by aversive stimuli and had weaker responses to salient threat and safety cues. On longer timescales, this gamma signal also correlated with increased skin conductance, a measure of autonomic state. Lastly, we found that direct electrical stimulation of the anterior insular cortex in a subset of subjects elicited self-reported increases in anxiety that were accompanied by enhanced frontoinsular beta oscillations. Together, these findings suggest that electrophysiologic representations of anxiety-related states and behaviors exist within anterior insular cortex. The findings also suggest the potential of reducing anterior insular beta activity as a therapeutic target for refractory anxiety-spectrum disorders.

## Introduction

Anxiety is a common symptom seen across numerous psychiatric disorders, and it is estimated that a third of individuals will develop one or more anxiety disorder in their lifetime [1, 2]. Current models of anxiety-spectrum disorders suggest that pathological anxiety arises from excessive, inappropriate activity within a distributed corticolimbic network that typically produces adaptive emotional responses to threats [3–5]. An improved understanding of neural mechanisms mediating prolonged elevated state anxiety could identify potential targets for therapeutic intervention.

Excessive anterior insular cortex activation has been observed across anxiety-spectrum disorders, including generalized anxiety disorder [6], panic disorders, social anxiety disorder [7], specific phobias [7–9], post-traumatic stress disorder [7, 8, 10], and obsessive-compulsive disorder [8, 11]. In all these disorders, insular activity is decreased by effective treatment [6, 11]. Functional neuroimaging data obtained in non-affected subjects have suggested that insular cortex also has a role in experimentally induced anxiety characterized by the anticipation and experience of perceived potential unpredictable threats, mirroring pathological insular activation seen across anxiety-spectrum disorders [12].

Here, we sought to identify intracranial electrophysiological biomarkers of prolonged state anxiety within the corticolimbic system, focusing in particular on anterior insular cortex given prior literature. Intracranial recordings allow for the precise identification of spectral-temporal neurophysiological changes that are not possible using noninvasive imaging modalities. We utilized the gold-standard translational paradigm for studying elevated state anxiety using a “threat of shock” protocol to experimentally induce temporary periods of elevated state anxiety [3, 13, 14]. This experimental approach is well-validated and elicits reliable subjective and physiological reports of anxiety that mirror states of pathological anxiety across disorders. We administered this task while simultaneously obtaining intracranial EEG recordings in medication-refractory epilepsy patient volunteers who were implanted with electrodes for purposes of surgical planning for treatment of their seizures. We then sought to identify associated neurophysiological correlates of reported anxiety, threat/safety cues, aversive stimuli, and autonomic state in the context of our induced anxiety task.

We found that anterior insula beta activity is selectively and persistently enhanced during periods of unpredictable aversive stimuli and that this beta activity is correlated with increases in state anxiety. In contrast, insular gamma was evoked by aversive stimuli and over longer timescales was correlated with skin conductance, a measure of autonomic outflow. Moreover, direct insular stimulation also increased self-reported anxiety and visceral sensations on a trial-to-trial bases, which associated with enhanced beta activity. Together, our results suggest that anterior insular cortex houses electrophysiological representations of anxiety-related states and behaviors.

## Results

### Self-reported Increases in Anxiety during Anxiety Induction Task

To identify neurophysiological correlates of subjective anxiety, we developed the anxiety induction task based upon the NIMH research domain criterion definition of ‘potential threat.’ The intent of the task was to activate brain systems for potential threat activated by the anticipation and experience of uncertain threat of aversive stimuli. This task consists of alternating epochs in which the participant is instructed by an auditory cue that they are in a ‘threat’ or ‘safe’ epoch (Fig 1A). Participants are told prior to the study that they can receive up to 10 electrical shocks to the fingers titrated to be irritating, but not frankly painful at random intervals during the threat epoch. Participants are also instructed that no aversive stimuli are delivered during the safe epoch. At the end of the task, subjects are asked to rate their anxiety level during the safe and threat epoch on a scale from 1 (‘no anxiety’ to 10 (‘worst anxiety ever’). Baseline trait anxiety ranged from minimal to severe as assessed by the Beck’s Anxiety Inventory (BAI) (Supplementary Table 1) administered prior to surgery.

**Figure 1:**
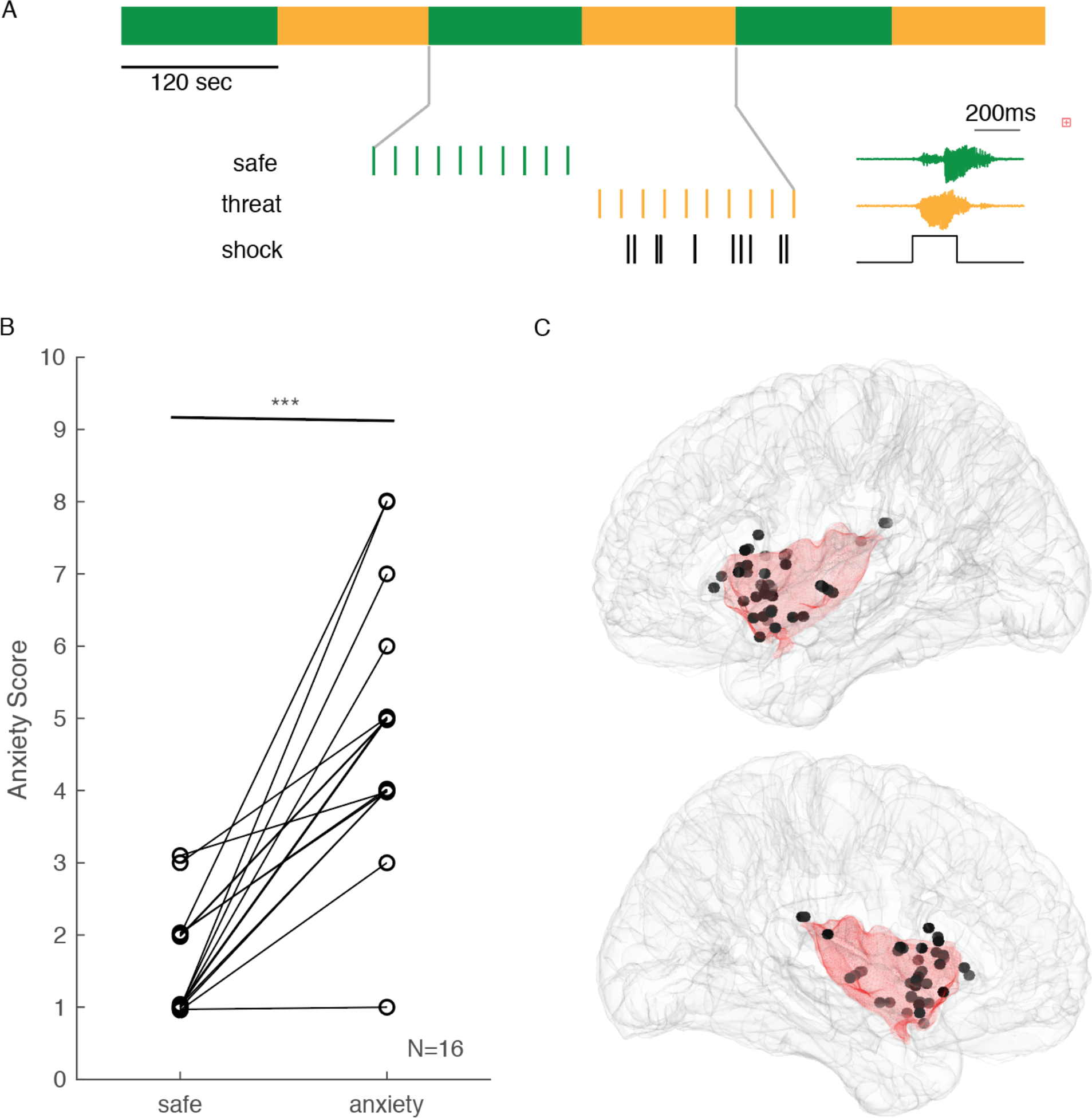
Intracranial Recording during Anxiety Induction Task: A) Stereoencephalography was recorded during the Anxiety Induction Task. This task involves alternating 2min safe and threat epochs. During Safe epochs, an auditory cue indicates to subjects that no aversive stimuli will be delivered. During Threat epochs, an auditory cue indicates to subjects that 10x aversive shock stimuli may be delivered at random interval during this period. B) Self-reported anxiety across Safe and Threat epochs during the Anxiety Induction task. C) Montreal Neurological Institute (MNI) template brain with coverage of recording electrodes (n=43) from 11 subjects in the insular cortex. *** p<0.0005, Wilcoxon signed-rank test

We identified sixteen subjects implanted with intracranial electrodes located within the corticolimbic network for purposes of identification of localizing their seizures based upon the assumption that these regions were likely to have anxiety-related activity. All subjects except one experienced an increase in reported anxiety during the threat epoch compared to the safe epoch (Fig 1B) (p=6.10x10^-5^, Wilcoxon signed rank). There was a trend for a relationship between task-induced changes in anxiety and pre-surgery BAI in our sample (r=0.52, p=0.084). Given prior literature implicating a role of the insula in anxiety, we sought to identify neurophysiological correlates of self-reported changes in anxiety within this region. Insular electrodes (n=43 electrodes from 11 subjects) were primarily located in the anterior aspect of the insula (Fig 1C).

Other corticolimbic regions which had electrode coverage included orbitofrontal cortex (n=70 electrodes from 9 participants), amygdala (n=23 electrodes from 8 subjects), and hippocampus (n=45 electrodes from 8 subjects), dorsal anterior cingulate (n=13 electrodes from 4 subjects), and inferior cingulate (n=7 electrodes from 4 subjects) (Supplemental Fig 1, Supplementary Table 2 for coverage). Intracranial recordings were performed in the context of the anxiety induction task. In 13/16 subjects, concurrent measures of skin conductance were collected during the task. We then sought to identify neurophysiological spectral changes associated with this increase in self-reported anxiety and task-related features within the insular cortex and other associated corticolimbic structures.

### Insular Beta Oscillations are Selectively Enhanced during Threat Epochs and Correlate with Anxiety

We performed spectral analysis of intracranial EEG data recorded across insular sites during the anxiety induction task and observed a persistent increase in beta (15-40Hz) activity with the transition from safe to threat epochs (Fig 2A). Aligning recordings to the transition between safe and threat epochs across electrodes, we observed a sustained increase in beta activity with the onset of the threat epoch across electrodes (Fig 2B).

**Figure 2:**
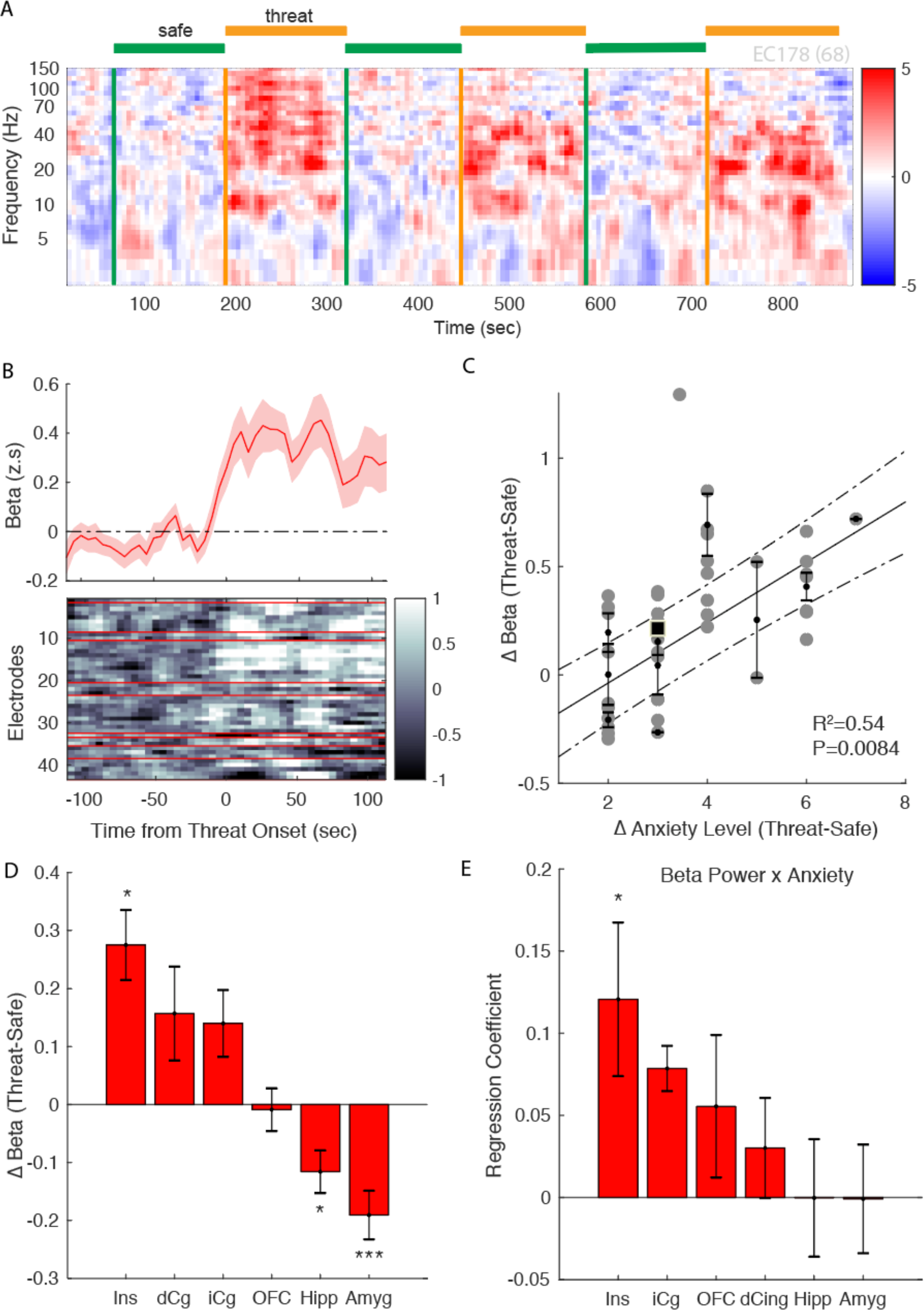
Insular Beta Correlates with Self-Reported Anxiety Across Task Epochs: A) Spectrogram of representative responses during the Anxiety Induction Task during Safe and Threat epochs. B) Average z-scored beta amplitude across all insular electrodes aligned to transitions from the Safe to Threat epoch (top panel). Shaded region represents SEM. Reponses across individual insular electrodes aligned to transition from safe to threat epochs. Electrode responses from subjects with larger changes in anxiety are depicted at the top (bottom panel). C) Change in beta amplitude correlated with change in self-reported anxiety during safe and threat epochs. Each dot represents average insular response across electrodes from a given subject with error bars representing standard deviation. Significance assessed using linear mixed effects model with epoch as covariate. D) Change in beta amplitude across corticolimbic structures. E) Regression coefficient between change in self-reported anxiety and beta amplitude across brain regions derived from a linear mixed effects model with change in anxiety as covariate. *p<0.05, *** p<0.0005

There was not a significant change in power in either insular theta or alpha (Supp Fig 2). The magnitude of the increase in beta activity between the threat and safe epochs was significantly correlated with the change in anxiety across epochs (R^2^=0.54, p<0.0084) (Fig 2C). This significant enhancement in beta activity was selective to the insula and was not observed in other corticolimbic regions such as the orbitofrontal, dorsal cingulate cortex, and inferior cingulate (Fig 2D). By contrast, beta activity was suppressed in the amygdala and hippocampus (Fig 2D). Importantly, only the anterior insular beta significantly predicted change in anxiety level during the task in contrast to beta activity in the dorsal cingulate cortex, inferior cingulate cortex, amygdala, orbitofrontal cortex, and hippocampus (Fig 2E). These data suggest that persistent anterior insular beta activity is a neural correlate of changes in state anxiety.

### Insular Beta Activity is Selectively Evoked by Cues Signaling Potential Threat

During the task, auditory threat and safety cues were presented to the subjects to notify them of whether they were in an epoch in which they might receive aversive stimuli at unpredictable times. Given that persistent increases in insular beta across epoch seemed to track with self-reported anxiety, we wanted to look at the temporal dynamics of the beta activity in relationship with task cues at a finer timescale. We found that insular beta activity was selectively enhanced by individual presentations of auditory threat cues, but did not respond to safety cues (Fig 3A,B). This increased beta activity was delayed occurring 2-5 seconds after the presentation of the auditory threat cue.

**Figure 3:**
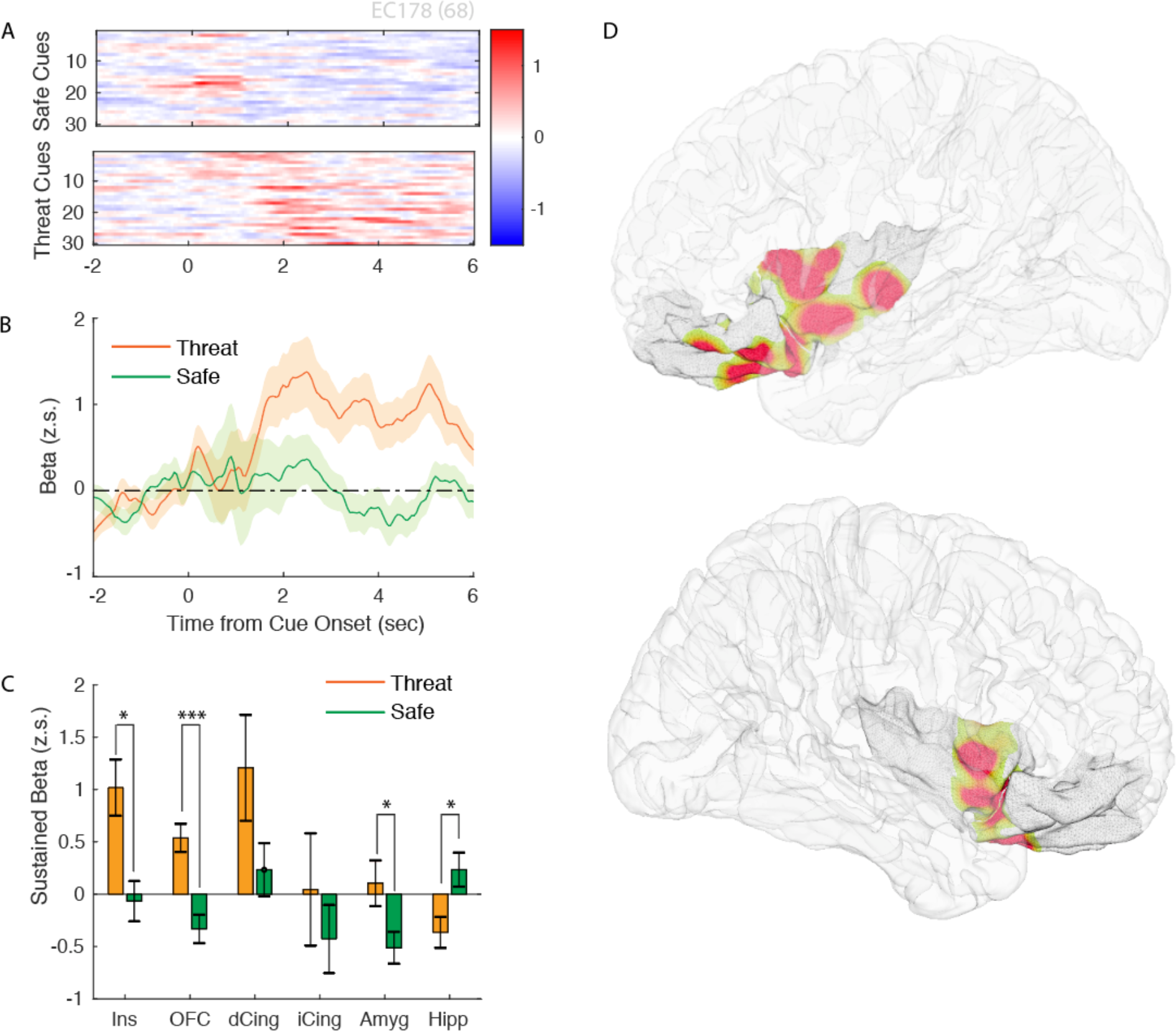
Fronto-Insular Beta is enhanced by Threat-Predictive Cues and Aversive Stimuli: A) Insular beta aligned to onset of auditory safe and threat cues across epochs. B) Average beta activity evoked by threat (orange) or safe (green) cues. C) Sustained beta (2-5 sec after cue onset) evoked by threat and safe cues across structures. Significance assessed using a signed rank test. D) Spatial distribution of evoked beta to threat vs safe cues within insular and orbitofrontal cortex. * p<0.05, ** p<0.005 , ***p<0.0005; Wilcoxon signed-rank test across electrodes.

Significant enhancement in sustained beta responses to threat versus safe cues was observed in orbitofrontal cortex (p=4.08x10^-5^, Wilcoxon signed rank test) in addition to anterior insular cortex (p=9.4 x10^-3^, signed rank test) (Fig 3C). Increases in dorsal anterior cingulate beta were observed but insignificant due to limited sampling.

Hippocampal beta showed the opposite response, significantly decreasing with auditory threat cues relative to safety cues (p=0.025, Wilcoxon signed rank test) (Fig 3C). Given that insular and orbitofrontal cortex demonstrated evoked beta responses to threat cues, we next sought to localize the anatomical distribution of these response. We found that the evoked beta activity at electrode contacts spanning the anterior insular cortex, frontoinsular junction, and posterior orbitofrontal cortex. Our limited coverage precludes us from exactly localize the spatial distribution of this activity, however. We also observed short-latency phasic increase in insular beta activity in response to aversive stimuli during the threat epoch (Supplemental Fig 3). These data suggest that the increases in anterior insula beta power across threat epochs are driven, in part, by beta activity evoked by cues signaling potential threat.

### Insular Gamma Responds More Robustly to Aversive Stimuli Compared to Other Task-Relevant Auditory Cues

In contrast to insular beta activity, short-latency, broadband insular gamma (30-150Hz) responses were evoked by all salient, task-relevant stimuli including aversive electrical stimuli, auditory threat cues, and safety cues. (Fig 4 A,B,C). At individual electrode sites, the insular gamma responses to aversive electrical shock stimuli was significantly greater than the responses to auditory threat (p=3.68x10^-4^, Wilcoxon signed rank test) and safety cues (p=2.00x10^-4^, Wilcoxon signed rank test) (Fig 4D). However, there was no consistent difference between the gamma response to auditory threat and safety cues (p=0.5144 ; sign-rank test). Responses to aversive stimuli were otherwise absent in other corticolimbic regions such as the OFC, inferior cingulate, amygdala, and hippocampus (Supp Fig 4). The insular gamma response to aversive stimuli was significantly greater than evoked responses in other limbic regions such as the OFC (p=1.43x10^-8^, rank-sum test), inferior cingulate (p=2.32x10^-4^, rank-sum test), amygdala (p=4.49x10^-6^, rank-sum test), and hippocampus (p=1.27x10^-9^, rank-sum test) (Fig 4E). In a more limited sample of 4 subjects with anterior cingulate coverage, we also observed gamma responses to aversive stimuli (n=8 electrodes, Supp Fig 4) that were not significantly different from the magnitude of anterior insular gamma responses (p=0.24, rank-sum test). These data suggest that anterior insular gamma selectively has robust short-latency responses to aversive stimuli in addition to more modest responses to task-relevant cues.

**Figure 4:**
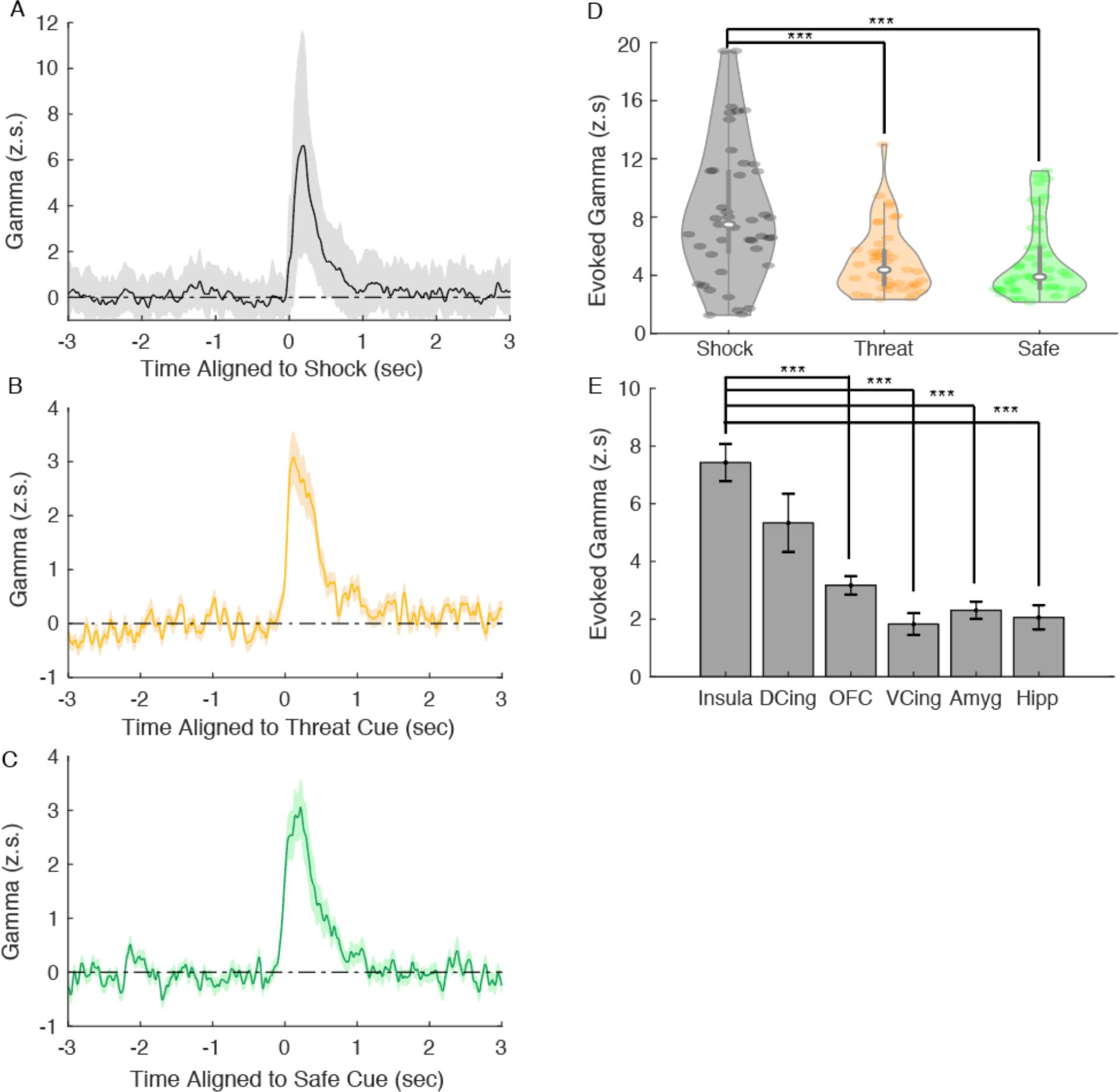
Insular Gamma Robustly Responds to Aversive Stimuli Insular gamma responses z-scored to baseline and aligned to A) aversive stimuli, B) auditory threat cue, and C) auditory safety cue. D) Evoked gamma responses at insular sites for aversive stimuli, threat cues, and safe cues. Significance determined with signed-rank test. E) Evoked gamma to aversive stimuli at insular, dorsal cingulate, orbitofrontal, ventral cingulate, amygdalar, and hippocampal sites. Significance determined by rank-sum test. ***p<0.0005

### Insular Gamma Activity Correlates with Skin Conductance Level

We also obtained skin conductance level measurements, a measure of sympathetic outflow [15, 16], in parallel with intracranial recordings in 13/16 subjects. In the remaining subjects, there was insufficient additional time to set up additional physiological sensors to obtain adequate quality recordings. In general, both the skin conductance level and insular gamma increased during threat epochs and decreased during safe epochs similar to insular beta activity (Fig 5A, C). Unlike beta activity, insular gamma activity was not significantly correlated to changes in anxiety associated during the task (p=0.2160 using linear mixed effects model). Instead, insular gamma was significantly positively correlated with skin conductance level in 8/11 subjects across recording sites (Fig 5B, Wilcoxon signed-rank test). In a more limited sample of 4 subjects with dorsal anterior cingulate coverage, we also observed that gamma correlated with skin conductance levels similar to the insular sites (n=12 electrodes, p=5.0x10^-4^ Wilcoxon signed-rank test, Fig 5B). This correlation was insignificant in other limbic regions such as the orbitofrontal cortex, inferior cingulate, amygdala, and hippocampus. By contrast, insular beta was not correlated with skin conductance level (p=0.83, sign-rank test). These data suggest that insular gamma on longer timescales monitors internal autonomic states.

**Figure 5:**
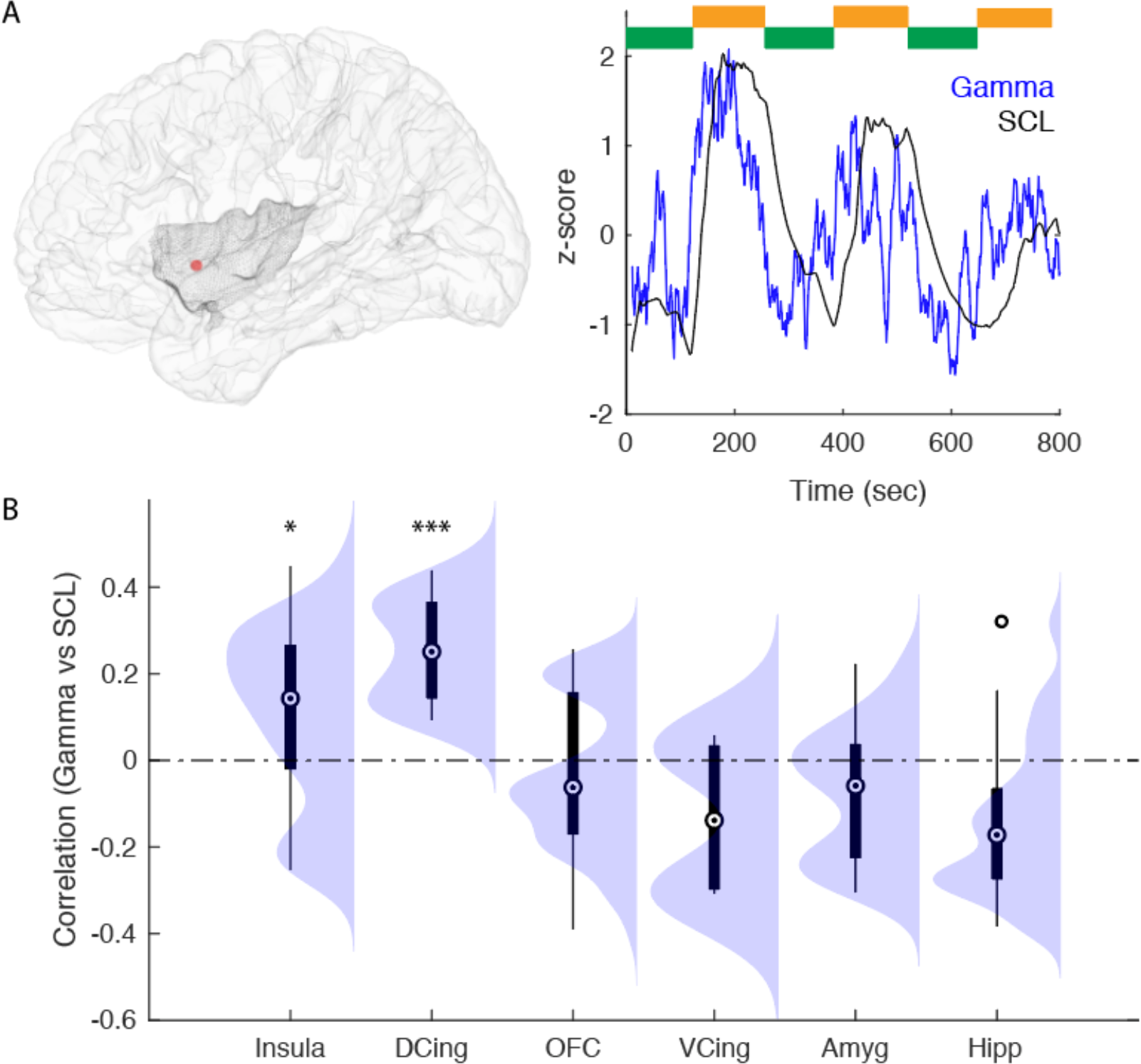
Insular Gamma Tracks Autonomic State. A) Left panel: Location of an example recording site (red) within insula (shaded). Right panel: Gamma activity recorded at example insular site (blue) with skin conductance level (black) across time. B) Spearman correlation between gamma activity across the insula, orbitofrontal cortex, amygdala, and hippocampus. *p<0.05,***P<5x10^-4^ significance assessed with signed- rank test.

### Direct Electrical Stimulation of Anterior Insula Increases Anxiety and Frontoinsular Beta activity

Our task-based intracranial recording results suggest a role for anterior insular beta activity in network activity associated with anxiety. We next sought to determine whether manipulating insular activity could generate changes in anxiety and related behaviors.

Stimulation studies were conducted in two participants within a window of time at the conclusion of their epilepsy monitoring unit inpatient stay after their seizures had been clinically localized prior to surgery. Direct electrical stimulation was delivered to the anterior insular cortex for brief trains of 5 seconds at 60Hz while recordings were conducted at neighboring frontoinsular sites at multiple levels of stimulation current intensity (Fig 6A). Anxiety scores were obtained prior to and during stimulation. In both subjects, there was a mild, but significant increase in anxiety during stimulation, which was only elicited at higher currents of 2-3mA and not at 1mA (Fig 6B, p=0.015 for EC205 and p=0.025 for EC206, paired t-test). This anxiety quickly dissipated upon the cessation of stimulation (Fig 6A,B- right panels). This increase in anxiety was accompanied by a visceral sensation such as a ‘rising feeling in the stomach’, ‘nausea’, ‘gut feeling that something is wrong,’ or a feeling ‘like someone I don’t like entered the room.’ We found that frontoinsular beta activity was significantly enhanced during stimulation on trials in both subjects (Fig 6A,C). In one subject (EC205), stimulation-induced beta was significantly greater at 2/3 electrode contacts during trials when subjects reported increases in anxiety than when they reported no change in anxiety level (contact FI5, p=0.0057 and FI6, p=0.0442, t-test). These increases were not found in neighboring ventromedial prefrontal contacts (Fig 6C), demonstrating the spatial specificity of these effects. These data suggest that direct electrical stimulation to the anterior insula enhances frontoinsular beta in addition to increasing anxiety associated with visceral sensations.

**Figure 6:**
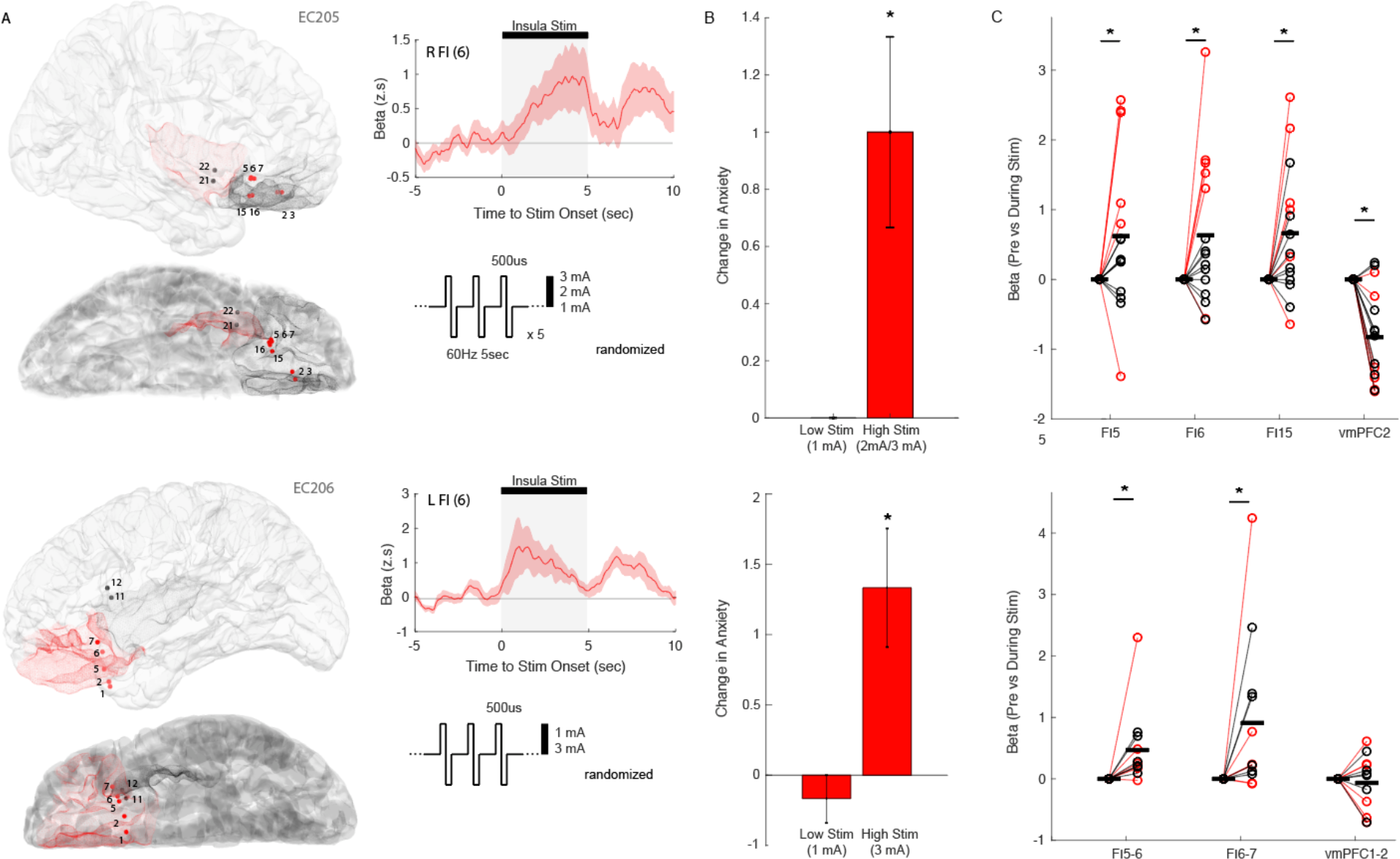
Insular Stimulation Can Elicit Self-reported Anxiety and Increases in Frontoinsular Beta Activity: A): Direct electrical stimulation of anterior insular cortex in 2 subjects. Bipolar sites of stimulation are depicted as black dot on brain and recordings sites are depicted as red dots. Low (1mA) or high (2/3mA) of 60Hz stimulation was delivered for 5 sec across multiple trials. Insular stimulation elicited an increase in frontoinsular beta activity. B): Insular stimulation elicited significant transient increases in self-reported anxiety during higher amplitudes of stimulation, but not at low amplitudes. C): Insular stimulation elicited beta activity at frontoinsular (FI) contacts but not neighboring vmPFC contacts across stimulation trials. Trials with an increase in anxiety are indicate by red lines and dots whereas trials with no changes in symptoms are indicated by black lines and dots. * p<0.05, paired t-test

## Discussion

Here, we show that insular cortex contains electrophysiologic representations of anxiety and associated behaviors across multiple timescales. Across subjects, beta oscillatory activity within the anterior insular cortex was persistently elevated during periods of uncertain aversive stimuli and correlated with state anxiety during epochs. This anxiety-related beta activity was specific to anterior insular cortex and not found in other corticolimbic structures. We observed that sustained beta activity was selectively evoked by cues signaling potential threat, but not safety cues. In contrast, we found that short-latency, phasic gamma was evoked by all salient stimuli within the task, but responses to aversive stimuli were the most robust and were largest in the insula compared to other corticolimbic sites. On a longer timescale, sustained insular gamma activity correlated with the skin conductance level, a measure of sympathetic outflow. Lastly, we also found that direct electrical stimulation of anterior insular elicited increases in anxiety in a dose-dependent manner that was associated with a self-reported visceral sensation and enhancement of frontoinsular beta activity.

These findings suggest a role for anterior insula in anxiety-related behaviors and generating states of anxiety.

These studies are consistent with prior functional neuroimaging studies using similar designs to experimentally induce anxiety, which identified prominent activation of anterior insular cortex [12, 17–19]. More broadly, the anterior insula has emerged as a critical component of an anxiety-related network that may function to represent the anticipation and experience of aversive stimuli [20–22], unpredictable threats [23], loss of perceived control [17], interoceptive states [24, 25], and prolonged levels of state anxiety. However, the ability to conduct direct electrophysiological recordings allowed us to observe all of these features as being simultaneously represented within distinct insular spectral bands across a range of timescales.

Multiple models exist to describe the role of the insula in anxiety. Amongst the first insights about the function of the human insula came from seminal work by Wilder Penfield, who identified that electrical stimulation of the anterior inferior insular cortex elicited a variety of visceral sensory responses [26]. This lead to early conceptualizations of the insula as being a part of the ‘visceral brain,’ receiving afferent interoceptive information from the body [27, 28]. Here, we have replicated Penfield’s initial insular stimulation findings and observe these visceral sensations are often accompanied by an increase in anxiety and modulation of beta activity within the fronto-insular cortex. Also consistent with a role for insular cortex in vicero-autonomic regulation, we find that insular gamma seemed to track with skin conductance level, a measure of sympathetic outflow. We previously demonstrated this relationship between insular gamma and skin conductance in a much more limited cohort in a virtual reality visual threat paradigm [29]. Subsequently, Craig has proposed a posterior-to-anterior progression in the insula whereby primary interoceptive signals are received in the posterior region while more abstract cognitive maps of bodily-emotional states are represented more anteriorly [28, 30]. In particular, imaging studies have identified the anterior insula as being a part of the ‘salience network,’ which functions to identify the most relevant external sensory information and internal emotional and bodily states to regulate cognition and behavior [31, 32].

Based upon our observations, we posit that anterior insular gamma represents salient external and internal stimuli to maintain a state of vigilance, responding phasically at short latencies to aversive stimuli, task-relevant stimuli, and interoceptive states. At a neurophysiological level, this broadband gamma activity may reflect the population activity of insular neurons similar to other regions of the brain [33, 34]. We also observe these gamma signals in other components of the salience network such as the dorsal anterior cingulate cortex. In contrast, insular beta cognitively filters and integrates this activity over a longer timescale, identifying cues predictive of potential threat, aversive stimuli, and other information to develop subjectively reportable states of anxiety. Perturbations of these insular representations, in turn, generate self-reported interoceptive sensations and increases in anxiety. In this way, insular beta and gamma activity provide a near complete representation of all task-relevant stimuli and measured anxiety-related behaviors in our experiments.

Our result that insular beta activity is a biomarker of state anxiety is consistent with other findings from our group. Previously, our group identified amygdala-hippocampal beta coherent activity over periods of tens of minutes as a biomarker of reported spontaneous changes in mood in medication refractory epilepsy patients with trait anxiety [35]. In our task, the epochs change every three minutes so we were not able to assess for this biomarker. Our other colleagues also have found that prefrontal beta spectral power recorded longitudinally from subdural leads correlating with ratings of depression and anxiety in Parkinson’s patients receiving DBS therapy [36]. Similarly, we observe strong orbitofrontal beta evoked response to threat cues in overlapping regions. Together, these studies suggest that organized beta oscillations across the corticolimbic system may serve to organize and generate states of subjective anxiety.

Our study has limitations. This study consisted of a relatively small sample of patients with medication-refractory epilepsy, which may not generalize to patients with primary anxiety-spectrum disorders or other populations. While we cannot rule out the possibility that the state anxiety biomarker that we identified has a seizure-specific etiology, the observed effects were robust across patients with a diversity of seizure pathologies. Also, recent studies have suggested that induced experimental induced anxiety and pathological anxiety both recruit activation of the insular cortex [12]. Our coverage also was limited in the corticolimbic network due to the fact that electrode placement was dictated by clinical need to localize seizures, and subjects had variable coverage. For this study, we only had electrode coverage over anterior insula so we have limited ability to comment on posterior insular functions. Also, subcortical regions such as the bed nucleus of the stria terminalis implicated in anxiety responses in this task were not able to be assessed here [18, 37]. Another caveat is that we only obtained changes in anxiety at the end of our task. However, retrospective reporting has previously been used in similar unpredictable shock tasks [5, 38]. Finally, it is unclear the extent to which the insular beta correlate of state anxiety that we identified under experimental conditions will hold in more ecologically valid environments. Future studies will need to test this possibility using longitudinal insular recordings in naturalistic settings, perhaps with chronic DBS implants.

Our identification of anterior insular beta oscillatory activity as a potential biomarker of state anxiety may be useful for targeting and optimization of non-invasive and intracranial therapeutic stimulation for treatment of anxiety-spectrum disorders. It also positions anterior insular beta activity as a potential biomarker for detecting pathological changes in state anxiety that might enable the development of “closed-loop” stimulation for delivering optimized treatment of severe, debilitating, treatment-resistant anxiety where the presence of elevated anterior insular beta activity is used to trigger intracranial stimulation only when needed.

## Methods

### Standard protocol approvals, registrations, and patient consents

The Institutional Review Boards of the University of California, San Francisco, and the University of Iowa approved the study protocol. Patients provided written informed consent prior to participation and all experiments were performed in accordance with the tenets of the Declaration of Helsinki.

### Participants

Participants were 16 neurosurgery patients with drug-resistant epilepsy undergoing invasive electrophysiological monitoring as part of their clinical care at the University of California, San Francisco. Depending on the clinical care objectives, participants were implanted sub-chronically with ECoG strips, grids, or SEEG depth electrodes. Additional participant information is provided in Supplementary Table.

### Electrophysiology acquisition and imaging

Intracranial electroencephalography (iEEG) recordings were obtained during quiet wakefulness. Local field potentials at each electrode were amplified and digitized according to the specifications in Table Table2.2. All electrodes were referenced to a subgaleal electrode. Each participant underwent a preoperative magnetic resonance imaging (MRI) session to acquire structural images of the brain. After intracranial electrode implantation, participants received a computed tomography (CT) scan, which was coregistered to the MRI for individual electrode localization. The Freesurfer anatomical atlas63 was used to localize each electrode to an anatomical region of interest (ROI), and for visualization purposes, individual participant data were warped to a common anatomical space (cvs_avg35_inMNI152)64.

### Anxiety Induction Task

This task is a modified version of the NIMH Research Domain Criterion’s suggested task for assessing the domain of potential threat. This task consists of alternating epochs in which the participant is instructed by an auditory cue that they are in a ‘threat’ or ‘safe’ epoch. Subjects are told prior to the study that they can receive up to 10 electrical shocks titrated to be irritating, but not frankly painful at random intervals during the threat epoch. No aversive stimuli are delivered during the safe epoch. Continuous measurements of skin conductance level were performed using electrodermal sensors placed on the first and second fingers.

Subjects were asked to rate their level of discomfort while individual electrical stimuli were uptitrated on a scale from 1 (‘no pain-barely able to feel stimuli’) to 10 (‘very painful’) with a 7 being the point where the stimulation is perceived as painful. The magnitude of the electrical stimulus was then adjusted to being a ‘6/10’. In most subjects, skin conductance and heart rate data was collected at the same time as intracranial recordings.

### Neural Recordings

We acquired electrophysiological recordings at a sampling rate of 3051.8 Hz using a 256-channel PZ2 amplifier or 512-channel PZ5 amplifier connected to an RZ2 digital acquisition system (Tucker-Davis Technologies, Alachua, FL, USA). Recordings for stimulation studies was conducted with the Natus EEG clinical recording system.

Signals were inspected across electrodes for epileptiform activity and electrical noise and if necessary, removed from further analysis. The local field potential was recorded from each electrode, notch-filtered at 60 Hz and harmonics (120 Hz and 180 Hz) to reduce line-noise related artifacts, and signals were downsampled to 400Hz. Electrodes were then re-referenced to the common average across channels sharing the same connector to the preamplifier (Cheung et al., 2016). For analyses involving task alignments, signals were bandpass filtered using the log-analytic amplitude of the Hilbert transform at 45 logarithmically-spaced center frequency bands within this range. The average analytic amplitude was then averaged in traditional EEG bands including theta (4-8Hz), alpha (8-12Hz), beta (15-40Hz), and gamma (40-150Hz) to generate spectrograms. Subsequently, this activity was binned in windows of 20sec shifted in to identify changes in activity aligned to safe and threat epochs or windows of 1sec shifted in 0.05 sec increments for responses to auditory safety/threat cues.

### Electrode Localization

Electrodes were localized by co-registering the preoperative T1 to a postoperative CT scan using in house software (Hamilton et al., 2017). Pial surface reconstructions were created from preoperative T1 MRI scans using Freesurfer. For visualization of electrode coordinates in MNI space, we performed nonlinear surface registration using a spherical sulcal-based alignment in Freesurfer, aligning to the cvs_avg35_inMNI152 template (Fischl et al., 1999). This nonlinear alignment ensures that electrodes on a gyrus in the participant’s native space remain on the same gyrus in the atlas space, but does not maintain the geometry of different electrodes.

### Statistical analyses

Group level analyses of response to safe/threat epoch and correlation with changes in self-reported anxiety were conducted using mixed effects linear regression with multiple power measurements within the same subject being represented as a fixed effects and variation in each observed recording being a random effects. We used the fitlme command in Matlab to generate regression coefficients and p-values.

## Acknowledgements

The authors are grateful to Maryam Bijanzadeh, Prasad Shirvalkar, Kristen Sellers, Ben Speidel, and members of the Chang Lab for helpful discussions during the course of this study. This research was funded by NIMH K23MH125018 and partially funded by the Defense Advanced Research Projects Agency (DARPA) under Cooperative Agreement Number W911NF-14-2-0043, issued by the Army Research Office contracting office in support of DARPA’S SUBNETS program. The views, opinions, and/or findings expressed are those of the author(s) and should not be interpreted as representing the official views or policies of the Department of Defense or the US Government.

## Author contributions

AML and E.F.C. designed the study. A.M.L. carried out the experiments, recruited subjects, performed the data analysis, and wrote the manuscript with input from other authors. E.F.C. supervised the experimental work.

