## Supplemental for "Human Anterior Insular Cortex Encodes Multiple Electrophysiological Representations of Anxiety-Related Behaviors"

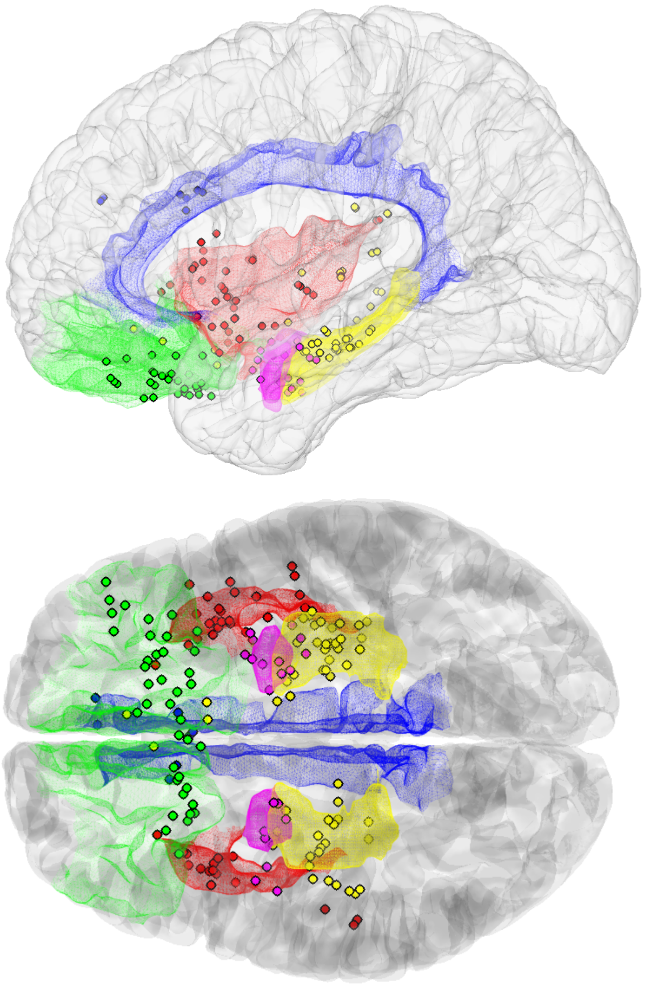

**Supplemental Fig 1: Coverage of Intracranial Electrodes**
Dots represent location of electrodes reconstructed in MNI space. Electrodes are color coated based upon region: red (insula), green (OFC), yellow (hippocampus), magenta (amygdala), cingulate (blue)

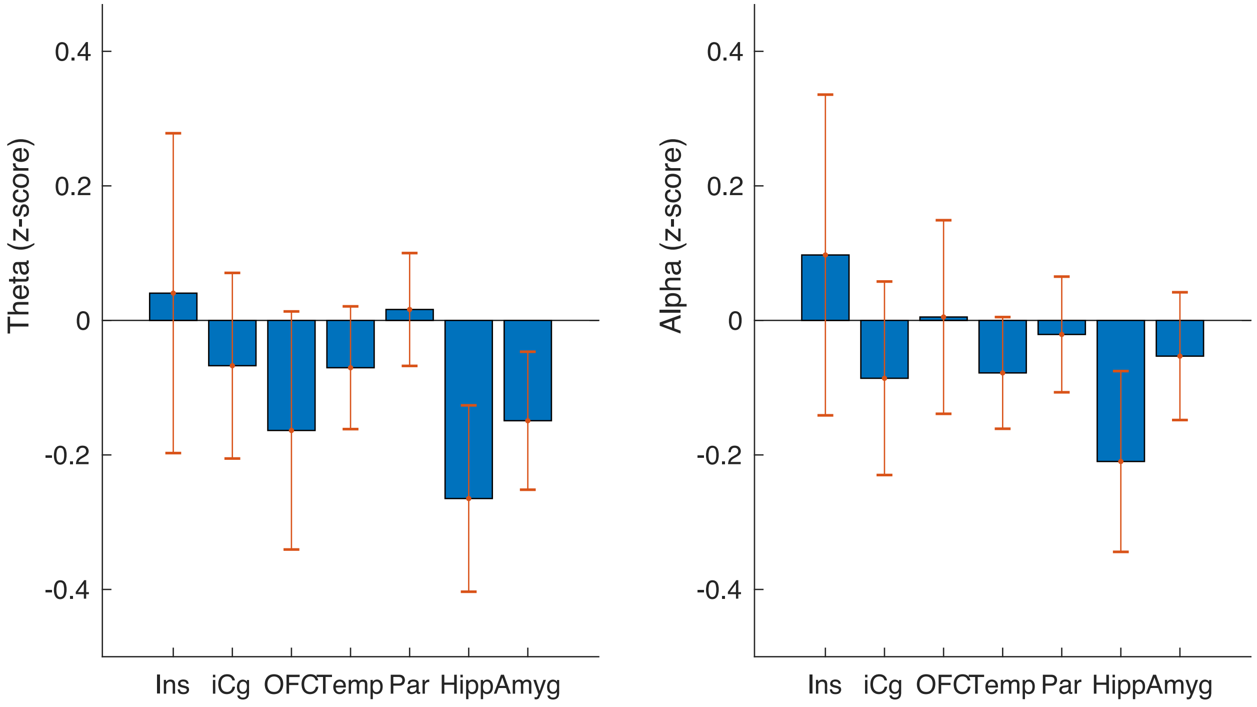


Supplemental Fig 2: No Change in Limbic Theta or Alpha during Threat Epochs

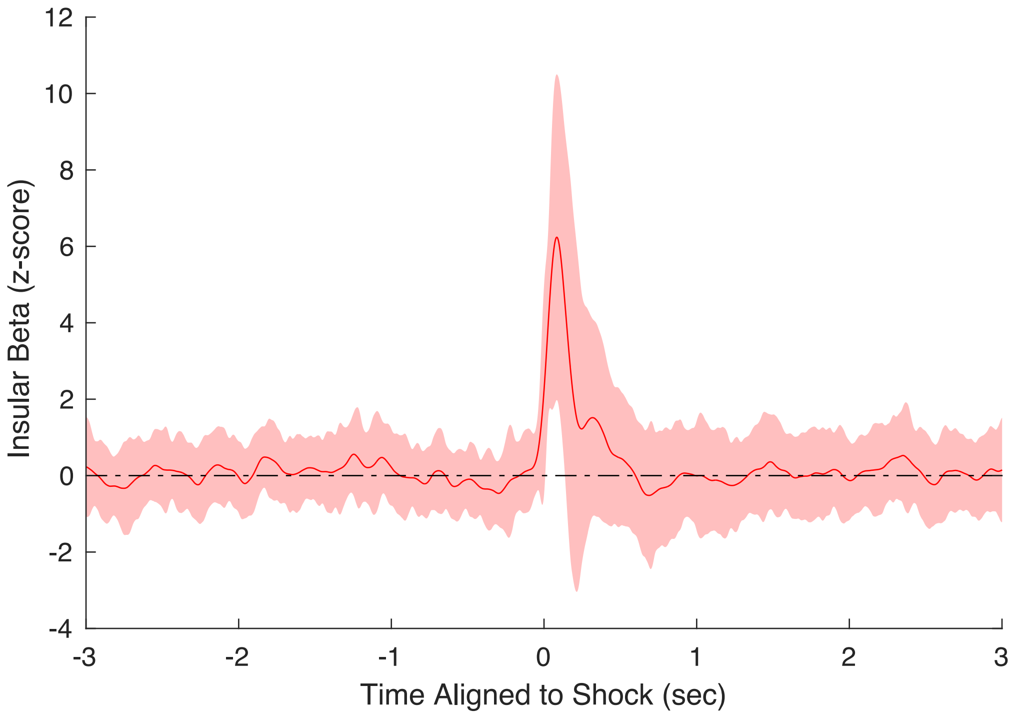


Supplemental Fig 3: Increase in Insular Beta in Response to Aversive Stimuli

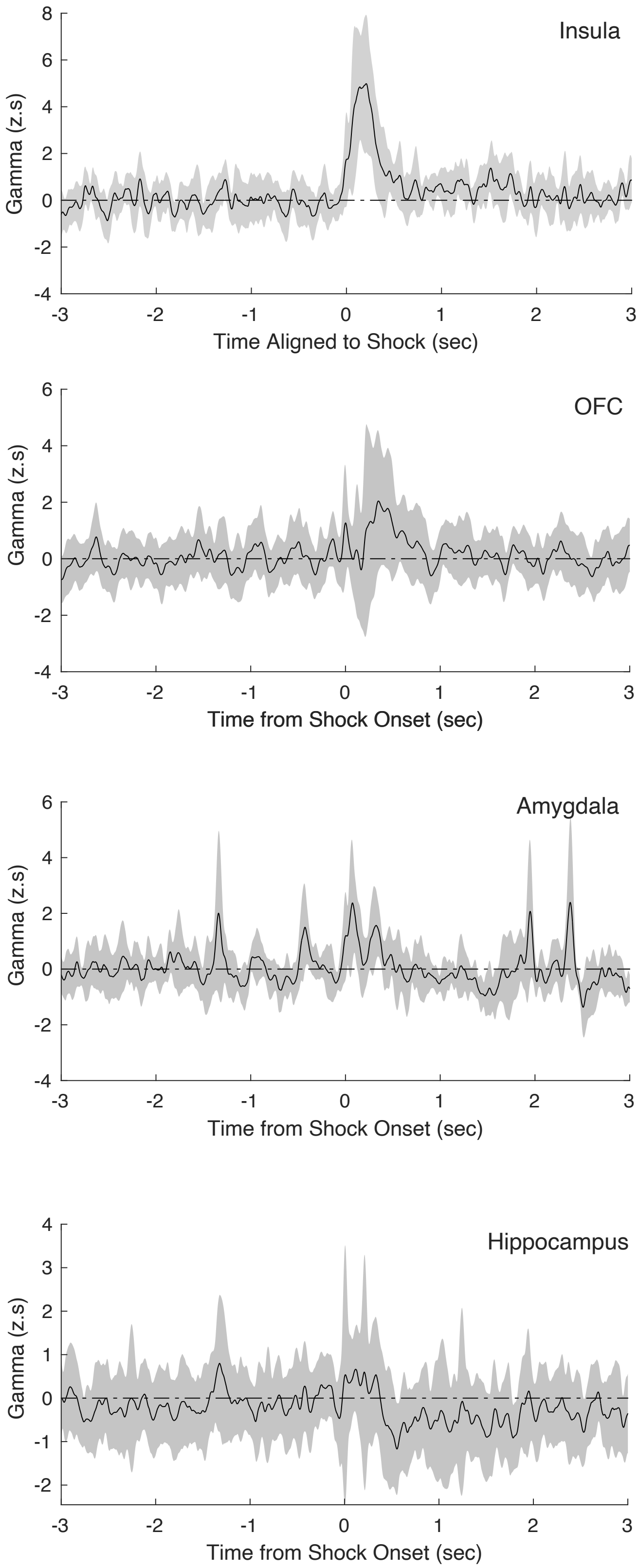


Supplemental Fig 4: Increase in Insular Gamma in Response to Aversive Stimuli, but Not in Other Limbic Structures

| | Anxiety Score | | | |
|---|---|---|---|---|
| Subject ID | Safe | Threat | BAI | BDI |
| EC150 | 2 | 8 | 26 | 35 |
| EC157 | 1 | 6 | 1 | 1 |
| EC158 | 1 | 7 | 43 | 28 |
| EC160 | 1 | 5 | | |
| EC162 | 1 | 4 | 10 | 4 |
| EC163 | 3 | 5 | 17 | 17 |
| EC178 | 1 | 5 | 0 | 0 |
| EC180 | 1 | 4 | 14 | 16 |
| EC181 | 2 | 7 | | |
| EC183 | 2 | 4 | 5 | 13 |
| EC184 | 1 | 8 | 39 | 38 |
| EC192 | 2 | 4 | 28 | 46 |
| EC200 | 1 | 1 | | 5 |
| EC201 | 3 | 4 | 10 | 3 |
| EC202 | 2 | 5 | 19 | |
| EC205 | 2 | 5 | 23 | 12 |
| EC206 | 1 | 3 | | |

**Table 1: Participant Self-Report for Anxiety Ratings and Beck's Anxiety Index (BAI) and Beck's Depression Index (BDI)**

| Region | N | Subjects | Electrodes |
|---|---|---|---|
| Insula | 11 | EC183,150,157,158,178,162,180,EC184,192,205,206 | 43 |
| OFC | 8 | EC150,158,178,162,183,200,201,206 | 39 |
| Inf Cingulate | 3 | EC150,162,184 | 7 |
| Amygdala | 8 | EC150,158,162,178,183,184,192,206 | 18 |
| Hippocampus | 8 | EC150,158,160,162,163,178,192,206 | 38 |

**Table 2: Electrode Coverage by Regions**
